# HLA class II polymorphism influences the immune response to protective antigen and susceptibility to *Bacillus anthracis*

**DOI:** 10.1101/841429

**Authors:** Stephanie Ascough, Rebecca J. Ingram, Karen K. Y. Chu, Stephen J. Moore, Theresa Gallagher, Hugh Dyson, Mehmet Doganay, Gökhan Metan, Yusuf Ozkul, Les Baillie, E. Diane Williamson, John H. Robinson, Bernard Maillere, Rosemary J. Boyton, Daniel M. Altmann

## Abstract

The causative agent of anthrax, *Bacillus anthracis*, evades the host immune response and establishes infection through the production of binary exotoxins composed of Protective Antigen (PA) and one of two subunits, lethal factor (LF) or edema factor (EF). The majority of vaccination strategies have focused upon the antibody response to the PA subunit. We have used a panel of humanised HLA class II transgenic mouse strains to define HLA-DR-restricted and HLA-DQ-restricted CD4+ T cell responses to the immunodominant epitopes of PA. This was correlated with the binding affinities of epitopes to HLA class II molecules, as well as the responses of two human cohorts: individuals vaccinated with the Anthrax Vaccine Precipitated (AVP) vaccine (which contains PA and trace amounts of LF), and patients recovering from cutaneous anthrax infections. The infected and vaccinated cohorts expressing different HLA types were found to make CD4+ T cell responses to multiple and diverse epitopes of PA. The effects of HLA polymorphism were explored using transgenic mouse lines, which demonstrated differential susceptibility, indicating that HLA-DR1 and HLA-DQ8 alleles conferred protective immunity relative to HLA-DR15, HLA-DR4 and HLA-DQ6. The HLA transgenics enabled a reductionist approach, allowing us to better define CD4+ T cell epitopes. Appreciating the effects of HLA polymorphism on the variability of responses to natural infection and vaccination will be vital in planning protective strategies against anthrax.

**Author Summary:** The bacterium responsible for causing the disease anthrax, Bacillus anthracis, produces a binary toxin composed of Protective Antigen (PA) and either Lethal Factor (LF) or Edema Factor (EF). Previous vaccination strategies have focused upon the antibody response to the PA subunit. However, within the field of bacterial immunity, there is a growing appreciation of the importance of the adaptive immune response, specifically led by CD4+ T cells. We identified long-term CD4+ T cell responses to PA epitopes following cutaneous human anthrax infection and vaccination, indicating that this toxin component is a principle B. anthracis antigen. To characterise the impact of polymorphism in HLA class II alleles at DR and DQ loci, we used transgenic mice to map the immunodominant epitopes from PA. This was correlated with survival in the transgenic lines following live anthrax challenge. We were able to demonstrate the differential impact of HLA class II alleles upon the CD4+ T cell immunodominant epitopes which shaped the immune hierarchy and therefore susceptibility to anthrax infection.

## Introduction

Anthrax is an acute zoonotic disease that primarily affects grazing mammals, although the causative agent, *Bacillus anthracis*, also infects humans and is found in many parts of the developing world, where the majority of natural human infection occurs [1]. Infections in humans, which may be fatal, depending upon the route of infection, are usually confined to agricultural workers, those who eat infected carcasses and those who handle the skins and coats of infected animals [2]. Over past decades, the need to protect individuals from occupational exposure has combined with fears regarding the use of anthrax as a bioweapon, to drive the development of vaccines based on the toxins produced by the bacteria [1]. Such concerns have resurfaced recently in relation to potential anthrax weaponisation [3]. Furthermore, there have been recent cases in Northern Europe of anthrax infections in intravenous drug users as a consequence of contaminated drug supplies [4]. There are also growing concerns regarding the effect of climate change in the Arctic upon the release of potentially viable anthrax spores from melting permafrost [5].

The three toxins of *B. anthracis*, Protective Antigen (PA), Lethal Factor (LF) and Edema Factor (EF) combine in a binary fashion, so that coupling PA with LF or EF produces Lethal Toxin (LT) or Edema Toxin (ET), respectively [6]. The two predominantly used vaccines, the United States-licensed Anthrax Vaccine Adsorbed (AVA; trade name BioThrax) and the United Kingdom-licensed vaccine, Anthrax Vaccine Precipitated (AVP), are culture filtrate vaccines containing PA and variable amounts of LF and EF [7]. Both vaccines are administered intramuscularly: AVA is given as three initial doses at 0, 1 and 6 months, while AVP is administered as a primary series of four vaccinations at 0, 3, 6 and 32 weeks [6]; a booster vaccination at 12 months, after the primary series for each vaccine, is then required. The requirement for an intensive vaccination regimen, as well as concerns about adverse reaction rates as high as 11% for the UK vaccine [8], and up to 60% for the US vaccine [9], have prompted interest in streamlined vaccination schedules or the development of effective, safe, subunit vaccines [10, 11].

Second-generation anthrax vaccines under development are based on the administration of the immunogenic anthrax toxins, specifically recombinant protective antigen (rPA). Human clinical trials have indicated that these rPA vaccines may be capable of eliciting robust cellular and humoral immune responses, whilst avoiding the adverse reactions associated with older filtrate-based vaccines [12-14].

PA-specific monoclonal antibodies generated from AVA-vaccinated humans were found to neutralise LT in vitro, and passive transfer of these antibodies provided protection in mouse models of LT challenge [15, 16]. Although it is possible to show passive transfer of immunity with toxin-neutralising antibodies [17], Crowe *et al.* found that over half of AVA-vaccinated individuals demonstrated no detectable toxin-neutralising effect; despite the presence of anti-PA antibodies in the majority of vaccinated individuals [18]. Studies in rhesus macaques have demonstrated that AVA administration is capable of providing protection from subsequent spore challenge, with a Th1/Th2 profile predictive of survival, even in the presence of very low levels of circulating anti-PA antibody [19].

Protection afforded by a response to PA in both rodent and non-human primate models has been suggested to be T-cell mediated [20, 21]. Plasmid vaccination in mice induces high antibody titres as well as PA-specific Th1 immunity and induction of a high level of IFNγ secretion [22]. Doolan and colleagues reported that individuals exposed to anthrax spores in the US mail service incident experienced dose-dependent priming of T cell immunity, and, to a lesser extent, of B cell immunity against PA [23]; low-level anthrax exposure led to PA T cell responses in the absence of detectable antibodies. While Glomski *et al* found that, in contrast to humoral immunity, IFNγ production by CD4+ T cells protected mice against capsulated B. anthracis infection [24].

Work from our lab has shown that individuals naturally exposed to anthrax spores demonstrate IFNγ secreting antigen-specific CD4^+^ T cell immunity to PA and LF, which for PA, showed correlation between the magnitude of response and the duration of the infection [25, 26]. We also found that a survivor of injectional anthrax developed strong, potentially protective, T cell immunity to several commonly immunodominant epitopes of PA and LF, previously described in Turkish patients [27]. This evidence suggests that cellular immunity has a critical role to play in vaccine mediated clearance of *B. anthracis*.

Whether the future of anthrax vaccinology lies with third-generation, subunit vaccines or with improved protocols for priming with existing vaccines, the need has never been greater to fully comprehend the nature of effective immunity to *B. anthracis,* and the impact of immunogenetic diversity. Here we describe a combined approach to characterising CD4^+^ T cell immunity to the PA toxin. This encompasses comprehensive analysis of T cell epitopes through investigation of HLA class II binding, mapping of responses in a panel of HLA class II transgenic mice, live challenge studies in HLA transgenic mice and studies of infected or vaccinated human donors. Our results show PA to be highly CD4+ T cell epitope-rich, with variable immunodominance which is dependent on HLA class II genotype. As discussed below, this has implications for wide-scale roll-out and assessment of PA-based vaccines.

## Results

### CD4+ T cell responses to B. anthracis PA epitopes in anthrax-recovered patients and vaccinees

We have previously described T cell memory responses to anthrax antigens in a cohort of individuals who suffered clinical disease after natural, occupational exposure [25, 26, 28]. These were agricultural workers from the Kayseri region of Turkey who had been in contact with infected livestock and been hospitalised with confirmed cutaneous anthrax infections. PBMCs were collected for immune analysis at 0.4 to 7.5 years after recovery under antibiotic therapy. In earlier studies, we described the fact that responses to recombinant PA and LF antigens were higher in naturally exposed individuals than in vaccinees receiving a full course of the UK AVP anthrax vaccine. Furthermore, immune responses in naturally infected donors were characterised by a broad cytokine profile, encompassing IL-2, IL-5, IL-9 and IL-13 [29]. In the present study we sought to analyse in greater detail the epitope specificity of vaccinated and infected individuals to PA. PA epitopes were screened by looking for ELISpot responses to a panel of 73 overlapping peptides of 20mers overlapping by 10 amino acid residues and analysed in pools of six. A total of 26 peptides were identified as epitopes in at least one AVP vaccinee (Fig 1), of which only 7 epitopes were an immune target for more than one vaccinee. Of note is the finding that only 4 vaccinees (AVP vaccinees donors 1-4) out of 10 responded to any epitopes, and of these the majority of the responses were elicited in donor 3, who responded to a total of 21 epitopes (Table S1). Although this study was not powered to make assumptions regarding the involvement of HLA alleles in the presentation of anthrax peptides, it is interesting that HLA-DR11 and DR13 were over-represented in the population of donors responding to the peptides contained within the vaccine. In contrast, the majority of infected individuals (7 out of 9 donors) responded to at least one PA epitope, and there did not appear to be any particular bias towards specific HLA alleles in the responses (Fig S2), with 69 of the 73 peptides analysed in this cohort found to carry infection-specific epitopes. Peptides such as PA 168-187 and PA 651-670 contained epitopes that were recognised with a high frequency response by multiple individuals (PA 168-187 mean = 264.2 spots/million, ±123.2 SEM, and PA 651-670 mean = 273.4 spots/million, ±123.6 SEM) and encompassing diverse HLA class II alleles. However, it is notable that although adjacent peptides (PA 161-187 and PA 641-660 respectively) were identified as epitopes for one of the vaccinated individuals, neither of the infection-specific epitopes, recognised in the context of multiple HLA alleles, were a focus of the response in any vaccinees.

**Figure 1.**
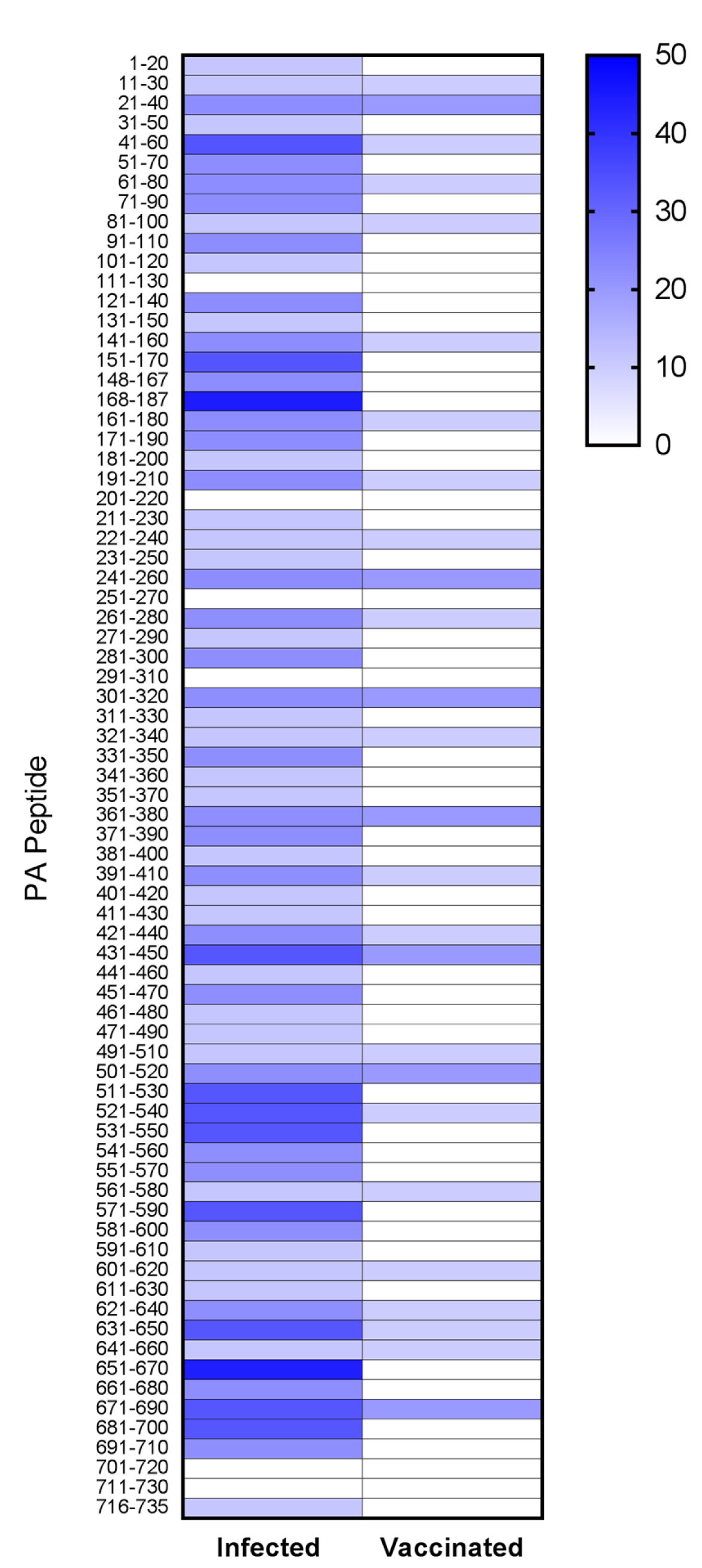
Heat map of CD4+ T cell epitope responses to anthrax PA domain I-IV peptides in human donors. Heat map representation of the epitope mapping results observed for positive CD4+ T cell IFNγ ELIspot responses in the human donor cohorts, comprising a total of 9 donors in the cutaneous anthrax (Kayseri) group and 10 donors in the AVP vaccinees (UK) group. Peptides were considered positive for the carriage of a CD4+ T cell epitope if the response was >50 SFC/10^6^ PBMCs and 2SD above negative control, and the stimulation index (peptide response/negative control response) value was ≥ 1.5. The domains were defined as described previously; domain 1 = PA 1-20 to PA 241-260; domain 2 = PA 251-270 to PA 471-490; domain 3 = PA 491-510 to PA 581-600; domain 4 = PA 591-610 to PA 716-735, with some peptides overlapping the boundaries between domains) [50]. The colour bar at the right indicates the percentage of donors responding to a given epitope, with shading from white (0%) to dark blue (50%).

In both infected and vaccinated cohorts, the epitopes came from sequences within all four domains of PA (Fig 1), indicating that, unlike LF, the majority of PA epitopes are not clustered within a single domain of the protein [25]. This comparison also highlights the fact that individuals who had been hyper-immunised on the standard UK schedule with seven to 14 doses of the AVP vaccine over 3.5 to 10 years, responded to fewer epitopes than infected individuals, with no epitopes identified that were present in the context of vaccination alone. This supported the suggestion, which we originally made in regard to LF; that live infection unveils cryptic anthrax epitopes not commonly recognised after administration of the protein antigen.

### Differential susceptibility to B. anthracis challenge in HLA transgenic mice

In order to more precisely define the contribution of different HLA class II alleles to anthrax and PA immunity, we turned to HLA class II transgenic mice as a defined, reductionist model allowing analysis of individual alleles in isolation.

We initially compared susceptibility of mice expressing either HLA-DR1 or HL-DQ8 to challenge with 1×10^6^ CFU (10^3^ median lethal doses, MLD) *B. anthracis* STI strain. HLA- DR1 mice were resistant to *B. anthracis* STI challenge (MLD > 10^6^ CFU), while HLA- DQ8 mice were also relatively resistant, with 80% survival. The more susceptible HLA class II transgenic mice demonstrated differential susceptibility to challenge at 10^5^ CFU (10^2^ MLD *B. anthracis* STI) with the following survival rates: DQ6 mice (100%), DR4 (80%), and DR15 (55%). By comparison, the parent strain for the HLA class II transgenics, C57BL6, showed 40% survival against a 10^5^ CFU contemporaneous challenge with the STI vaccine strain of *B. anthracis*.

The bacterial loads recovered from the spleens of individual surviving mice of each strain at day 20 are shown in Fig 2. In general the mean bacterial loads in spleens at day 20 post-infection were lower than, but proportional to, the original challenge dose level. The groups challenged with 10^6^ CFU (DR1, DQ8) had high bacterial loads, although the mean bacterial loads for the DQ6 mice (challenged with 10^5^ CFU) did not differ significantly from those for the DR1 or DQ8 mice, which had been challenged with ten-fold more bacteria, suggesting that the DQ6 mice were slower to clear the infection.

**Figure 2.**
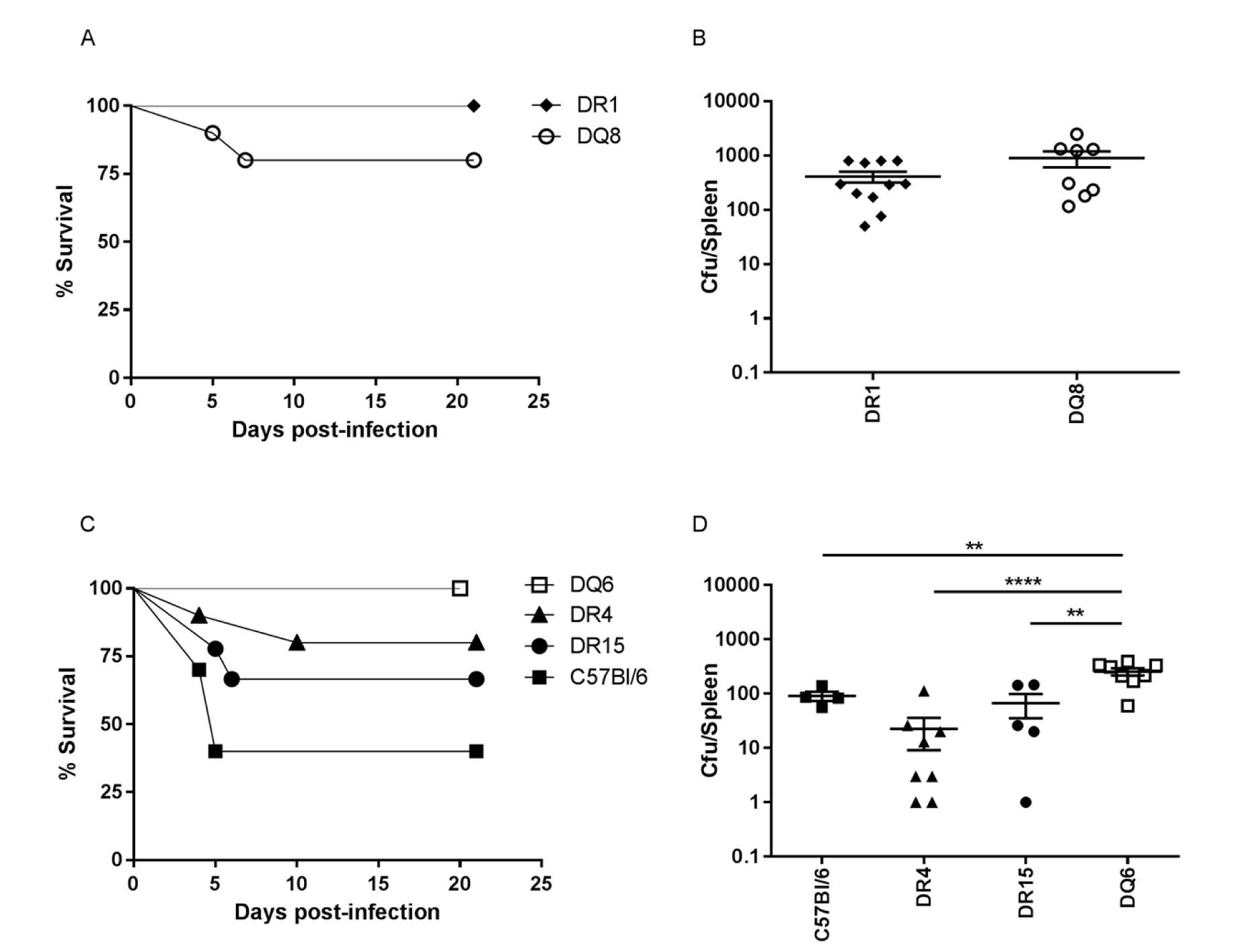
Differential susceptibility of HLA class II transgenic mice to anthrax infection. Groups of naïve HLA transgenic (DR1 n=11, DQ8 n=10, DR15 n=9, DR4 n=10, DQ6 n=8) or C57Bl6 (n=10) mice were challenged with either 10^5^ (C and D) or 10^6^ (A and B) CFU *B. anthracis* STI strain, in order to compare susceptibility. Mice were challenged intraperitoneally and their survival observed for 20 days post-infection. Percentage survival, together with mean splenic bacterial counts per HLA type, is shown for mice succumbing within the observation period (days 1 to 19) and for survivors culled at day 20. Statistical comparison of mean bacterial loads by mouse strain (D) indicated that higher bacterial loads were seen in DQ6 in comparison to; C57BL/6 (** p=0.0093), DR4 (**** p<0.0001) and DR15 (** p=0.0014), (One-way ANOVA, Tukey’s multiple comparisons).

HLA transgenic mice were less susceptible to infection with *B. anthracis* STI strain than the parent strain C57BL6 mice. HLA-DR1 mice were resistant to infection with a high-level challenge (10^6^ CFU). DQ6 strain mice were resistant to 10^5^ CFU and relatively slow to clear the infection. The order of susceptibility of mouse strains to *B. anthracis* infection was determined to be: C57Bl6 > DR15 > DR4 > DQ6 > DQ8 > DR1.

### CD4^+^ T cell responses to B. anthracis PA epitopes in HLA transgenic mice

The greater immunogenetic complexity of HLA-outbred human populations makes it considerably more challenging to define the restricting HLA molecule responsible for antigen presentation; the HLA class II transgenic mouse models offer a reductionist system in which to define HLA-restricted epitopes of relevance to humans carrying the same alleles. Using these transgenic models in protein and peptide immunisation we were able to build a comprehensive picture of immunodominant HLA class II restricted epitopes derived from PA. Mice were immunised with the recombinant PA protein and draining lymph node cells were restimulated with a peptide library spanning the PA sequence (73 peptides in total, with some peptides overlapping the boundaries between domains: domain 1 = PA 1-20 to PA 241-260; domain 2 = PA 251-270 to PA 471-490; domain 3 = PA 491-510 to PA 581-600; domain 4 = PA 591-610 to PA 716-735,). After immunisation with the recombinant protein of interest, all HLA transgenic mice responded to the whole rPA (Fig 3), but the response to the individual peptides was found to be HLA-specific.

**Figure 3.**
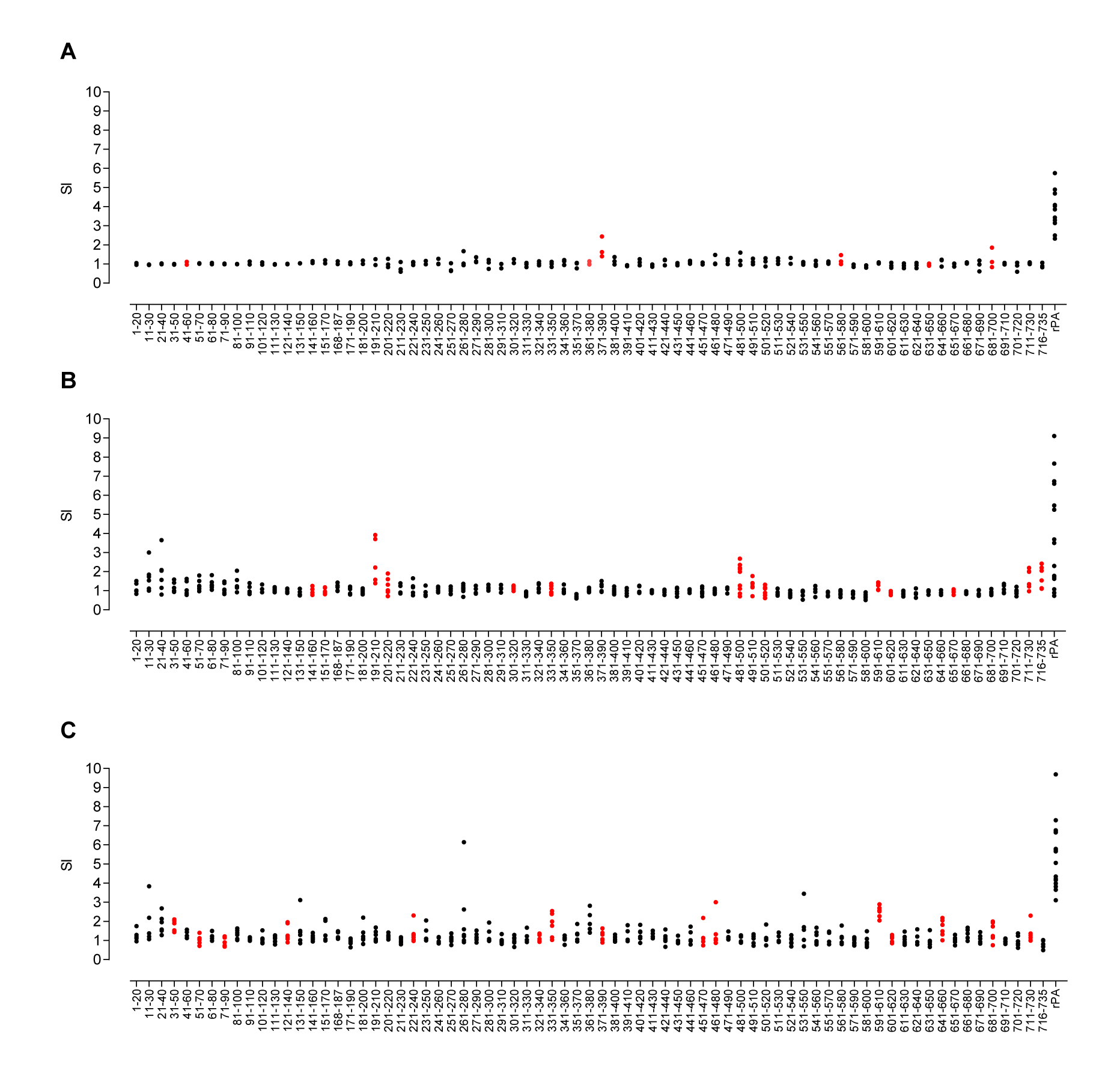
T-cell responses to PA peptides in whole rPA-immunised HLA-DR and HLA-DQ transgenic mice. Groups of HLA transgenic mice were immunised with the whole rPA protein in adjuvant, and the proliferative responses of draining lymph node cells to overlapping synthetic peptides representing the complete PA sequence were determined. Scatter plots show responses of individual mice transgenic for (A) HLA-DR1 (n=3 for each peptide data point, and n=11 for the rPA data point), B) HLA-DQ8 (n=6 for each peptide data point, n=18 for the rPA data point) and C) HLA-DR4 (n=6 for each peptide data point, and n=17 for the rPA data point). Data is presented as the SI calculated as the mean CPM of triplicate wells in the presence of peptide divided by the mean CPM in the absence of antigen. Values twice the mean CPM in the absence of antigen were considered positive responses. Confirmed epitopes are highlighted in red.

We investigated whether there might be any correlation between susceptibility of the HLA transgenic lines to challenge and the breadth of T cell epitope recognition. Antigen-specific T cell responses to all stimulatory peptides were further investigated by peptide immunisation and screening (Figs S1, S2 and S3). In total, 6 HLA-DR1 restricted epitopes were identified: PA 41-60, PA 361-380, PA 371-390, PA 561-580, PA 631-650, and PA 681-700 (Fig 3A and Fig S3). In comparison 14 HLA-DQ8 restricted epitopes were identified: PA 141-160, PA 151-170, PA 191-210, PA 201-220, PA 301-320, PA 331-350, PA 481-500, PA 491-510, PA 501-520, PA 591-610, PA 601-620, PA 651-670, PA 711-730, and PA 716-735 (Fig 3B and Fig S1): and 15 HLA-DR4 restricted epitopes were identified: PA 31-50, PA 51-70, PA 71-90, PA 121-140, PA 221-240, PA 321-340, PA 331-350, PA 371-390, PA 451-470, PA 461-480, PA 591-610, PA 601-620, PA 641-660, PA 681-700, and PA 711-730 (Fig 3C and Fig S2). Whilst some of these epitopes were recognised by more than one HLA type (PA 331-350, PA 591-610, PA 601-620 and PA 711-730 were constituents of both DR4 and DQ8 responses, while PA 371-390 and PA 681-700 were recognised by both DR1 and DR4 alleles), no one epitope was found to provoke a response in all 3 HLA alleles tested. Thus, it was noteworthy that HLA-DR1 transgenic mice, which were the least susceptible to anthrax challenge, responded to fewer epitopes with a reduced repertoire of CD4+ T cell recognition than the other HLA alleles screened.

### The differential PA peptide binding across distinct HLA polymorphisms

Overlapping 20-mer peptides that represented the whole PA protein sequence were evaluated for binding affinity to seven common HLA-DR alleles and two common HLA-DQ alleles (Table 1). The two epitopes that were recognised by multiple individuals from the infected cohort (PA 168-187 and PA 651-670) showed a complete disparity in their HLA binding affinities. Whilst PA 168-187 was not recognised by any of the transgenic lines and showed an exceptionally low binding affinity across all HLA-DR alleles tested, PA 651-670 showed strong-to-moderate binding across all HLA-DR alleles, and bound strongly to HLA-DQ8, which also correlated with a strong response seen in the corresponding transgenic line. Overall, we were not able to identify a propensity towards a strong HLA binding affinity in those epitopes that were a feature of the infected response. In contrast, all but one (PA 501-520) of the seven epitopes identified in more than 20% of the vaccinated cohort demonstrated high binding affinities for the HLA-DR or DQ alleles carried by those individuals. This suggests that the binding affinity may be a more important predictor of epitope hierarchy in the context of vaccination than infection.

**Table 1.**
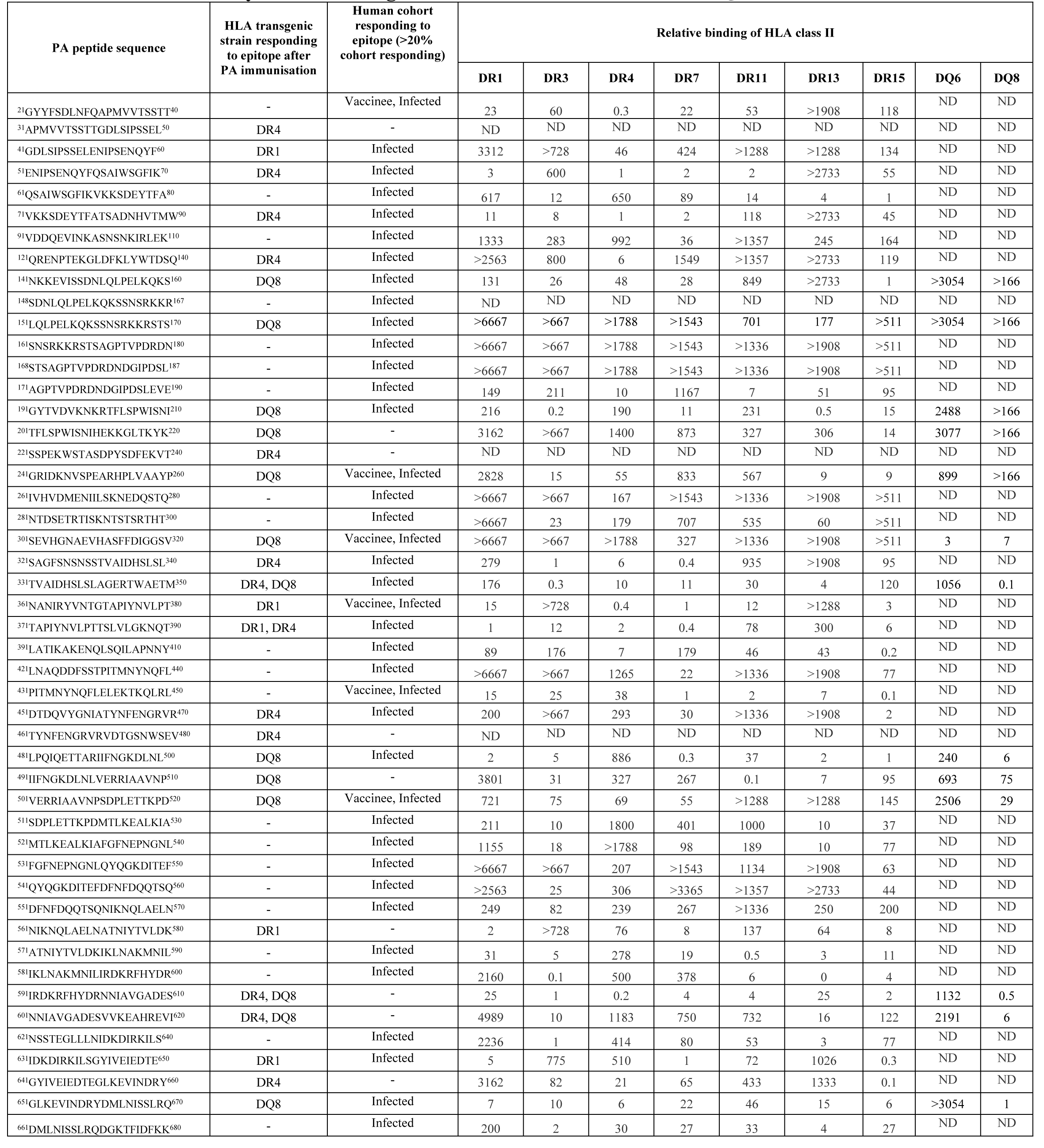
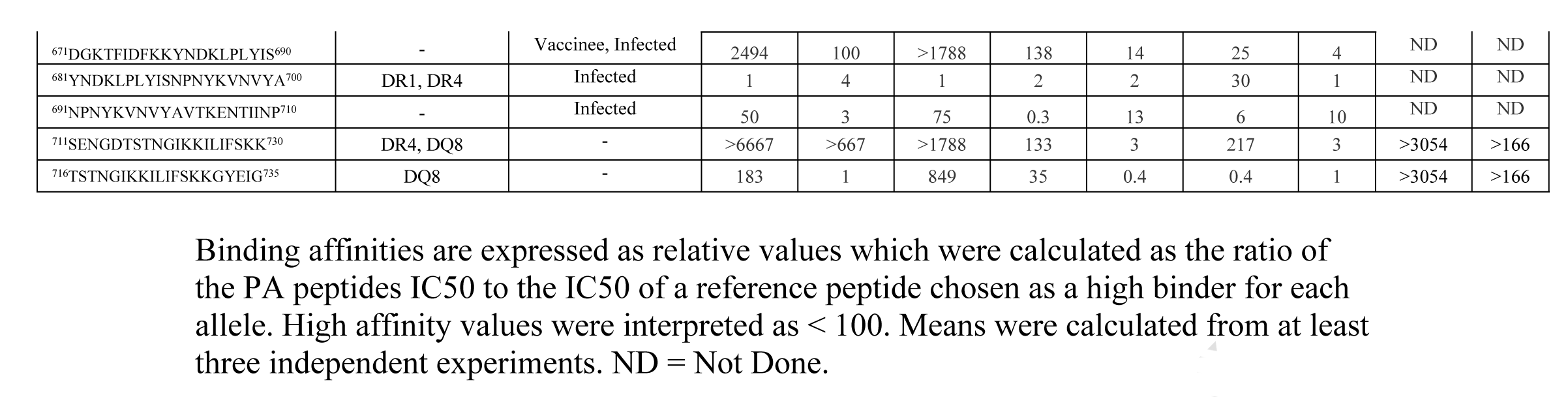
The PA peptides, identified in transgenic mouse strains and human cohorts, show relatively broad binding to common HLA-DR and HLA-DQ alleles.

## Discussion

Human exposure to anthrax spores continues to be of considerable concern in diverse spheres of clinical infectious disease; most commonly, exposure may occur naturally, either after ingestion of infected animals or through contact with infected animal products. Other routes of exposure could occur through deliberate release, acts of bioterrorism, or injection of contaminated drugs by intravenous drug users [3, 27, 30]; in these contexts, especially the threat of bioterrorist use, there has long been a perceived need to have an effective anthrax vaccination programme available. Three major vaccines have been in use in various parts of the world since the Cold War, with various recombinant subunit vaccine candidates in trial for rollout [31, 32]. Interestingly however, compared to many other bacterial pathogens, the immunology and immunogenetics underpinning any clear understanding of correlates of protection (CoP) are poorly delineated for anthrax [33]. Although vaccine development has focused largely on the endpoint of PA-targeted neutralising antibody, this alone is unlikely to confer sterilising immunity. At a general level, the CoP for effective AVA-vaccine-induced protection of macaques from anthrax challenge are IgG titre and IFNγ^+^ T cell frequency against PA [34]. Protection conferred by anthrax spores is entirely dependent on CD4+ T cells [35].

In seeking an improved understanding of the interaction between *B. anthracis* and protection by the human immune system, a key question has been the impact of immunogenetic heterogeneity at the population level [36]; work in mouse models has suggested that, as expected, both MHC and non-MHC polymorphisms are influencing these factors [37]. With respect to human vaccination, there is evidence of reduced immune responsiveness to PA in individuals with the DRB1*1501/DQB1*0602 haplotype [38]. In light of the importance of anti-PA immunity for protection and the relatively high frequency of this haplotype in many human populations, there is cause for concern in relation to vaccine efficacy and vaccine confidence. The situation is reminiscent of hepatitis B virus and MMR vaccinations, both of which demonstrate the profound influence of HLA polymorphism [39, 40].

Our aim here has been to shed light on the role of HLA class II alleles in PA epitope presentation to the immune system and thus on disease outcome after anthrax challenge. A key experiment in this regard was to compare the impact of STI challenge on survival and the control of bacterial load in mice, all on a C57BL/6 background and lacking expression of endogenous murine MHC class II heterodimers but differing in expression of specific HLA-DR or HLA-DQ alleles. The background C57BL/6 strain is considered one that mounts a low antibody response to anthrax PA and LF [37]. We found that HLA-DR15 transgenics (expressing the HLA-DRB1*1501 allele) were the most susceptible to challenge, echoing the results of human AVA HLA-DRB1*1501^+^ vaccinees [38]. It is particularly noteworthy that the effects of HLA class II alleles must be differentially effective in CD4+ T cell-mediated control of bacterial dissemination during the first 4 to 6 days after challenge, the very earliest days of detectable priming of an adaptive immune response. Nuanced differences in the potency and frequency of the initial CD4+ T cell responses have the potential to favourably impact survival by driving cellular responses to intracellular infection and generation of an initial neutralising antibody response. Such differences in susceptibility due to HLA polymorphisms are unlikely to have imposed evolutionary selection pressure in anthrax-exposed human populations. The pathogen is rarely transmitted from human-to-human, outbreaks tend to be of a limited nature (such as a local community consuming the same contaminated livestock), and most cases are not fatal. The greater concern relates to potential gaps in the efficacy of large-scale vaccination programmes for biodefense purposes, such as in the US military.

We looked at mechanisms underpinning HLA differences in susceptibility, starting with mapping of CD4+ T cell epitopes from PA. Our key findings were that natural infection elicits a considerably broader CD4+ epitope response than AVP vaccination and at least in the setting of natural infection, this is a very epitope-rich sequence, with epitopes spanning the entire length of the protein. It is well-established that in communities where environmental exposure to anthrax is relatively common, such as among goat-herders, symptomatic exposure confers lifelong protection from re-infection [25]. Differences in antigen processing and generation of epitopes for HLA class II binding between the AVP subunit vaccine components and live infection of APC might in some respects have been predictable, except that earlier studies of dendritic cells treated with lethal toxin showed a complete loss of the ability to effectively stimulate peptide-specific CD4+ T cells [41]. The PA sequence contains a number of regions with potential broad-ranging immunogenicity in terms of high-affinity binding to the majority of HLA class II alleles tested: 5 of the PA peptides analysed are relatively unusual in their capacity to bind very diverse HLA class II heterodimers at high affinity; PA191-210, 331-350, 481-500, 591-610 and 711-730. The 191-200 PA epitope overlaps one that we have previously identified at the CD4+ T cell level as being strongly recognised in the memory T cell response of a 60-year old intravenous drug-user who survived injection of anthrax-contaminated heroin [27]. This collection of epitopes would be excellent candidates for a highly immunogenic, widely applicable, epitope-string vaccine. Importantly, the fact that all bind HLA-DRB1*1501 with high or very high affinity makes it likely that the ‘low-responder’ status of HLA-DRB1*1501 vaccinees would be overcome by an approach focused on these epitopes. However, HLA class II-related differences in susceptibility to anthrax challenge cannot be a simple question of relative availability of high-affinity HLA class II-binding PA epitopes to activate the CD4+ T cell repertoire: the most susceptible HLA allele that we identified, HLA-DRB1*1501, can present at least as many PA epitopes as can the least susceptible allele, HLA-DRB1*0101. It is also important to stress that, while the HLA transgenic mice used to define immunodominant PA epitopes offer a useful reductionist system, the immune responses seen in this system may not fully recapitulate the effect of the individual HLA polymorphisms in a complete immune system. This may give a partial explanation for the divergence in epitopes identified in the HLA transgenics and those found in the human cohorts.

In summary, we draw two important conclusions from this comprehensive analysis of T cell recognition of anthrax PA. The first is that PA is an unexpectedly epitope-rich antigen, whether considered from a perspective of HLA class II binding or of CD4+ T cell recognition. The second key point, and one that offers an important note of caution to vaccinologists and to those planning biodefense strategies, is that there are likely to be major differences in both vaccine efficacy and anthrax severity imposed by HLA polymorphism within the population. These factors underscore the importance of considering immunological and vaccination strategies that can overcome such differences.

## Materials and Methods

### Ethics Statement

Human blood samples from Kayseri (Turkey) were obtained with full review and approval by The Ethics Committee of the Faculty of Medicine, Erciyes University. Human vaccinees based at DSTL, Porton Down, participated in the context of a study protocol approved by the CBD IEC (Chemical and Biological Defence Independent Ethics Committee). Written informed consent was obtained from all human volunteers. All mouse experiments were performed under the control of UK Home Office legislation in accordance with the terms of the Project License (70/5994) granted for this work under the Animals (Scientific Procedures) Act 1986, having also received formal approval of the document through the Imperial College Ethical Review Process (ERP) Committee.

### HLA class II transgenic mice

HLA class II transgenic mice carrying genomic constructs for HLA-DRA1*0101/HLA- DRB1*0101 (HLA-DR1), HLA-DRA1*0101/HLA-DRB1*0401 (HLA-DR4), HLA- DRA1*0101/HLA-DRB1*1501 (HLA-DR15) and HLA-DQA1*0301-DQB1*0302 (HLA-DQ8), crossed for more than six generations to C57BL/6 H2-Ab^00^ mice, were as described previously [42-46]. All experiments were performed in accordance with the Animals (Scientific Procedures) Act 1986 and were approved by local ethical review panel.

### Live B. anthracis challenge

Preliminary data indicated that there was a divergence in the susceptibility of mouse strains to anthrax challenge. Therefore, naïve mice were challenged with *B. anthracis* STI strain by the intraperitoneal route at one of two dose levels: 11 HLA-DR1 and 10 HLA- DQ8 mice were challenged with 10^6^ colony forming units (CFU) while 9 HLA-DR15, 10 HLA-DR4, 8 HLA-DQ6 and 10 C57Bl6 were challenged with 10^4^ CFU per mouse. The animals were monitored for 20 days post-infection, after which all survivors were sacrificed and their spleens were removed and homogenised in 1 mL of PBS before plating out onto L-agar plates. Colonies were counted after 24 hours of culture at 37°C, and the mean bacterial count per spleen was determined.

### Expression and purification of PA antigens

Good Manufacturing Practice grade rPA was provided by Avecia Vaccines (Billingham, UK) and had endotoxin levels of < 1 EU/mg. Individual domains of PA and peptides were expressed in E. coli and purified as previously described [47]. All proteins and peptides were resuspended in DMSO at 25 mg/mL.

### PA epitope mapping in transgenic mice

Mice were immunised in one hind footpad with 50 μL of 12.5 μg recombinant full-length PA, PA peptides, or a control of PBS, emulsified in an equal volume of TiterMax Gold (Sigma-Aldrich, USA) by syringe extrusion. After 10 days, immunised draining popliteal lymph nodes were removed and disaggregated into single-cell suspensions by filtration through 0.7 μm cell strainers. Lymph node cell responses were recalled *in vitro* with 25 µg/mL of either rPA, truncated PA domains comprising the PA protein, or the overlapping 20-mer peptides covering the full-length PA sequence. This produced a CD4+ T cell epitope map of the entire PA protein sequence. To confirm the immunodominant epitopes identified by this large-scale mapping, mice were then immunised subcutaneously with 12.5 μg of the individual PA peptides in TitreMax adjuvant. After 10 days the lymph node cells were challenged *in vitro* with 25 µg/mL of the recombinant full-length PA and the immunising and two flanking PA peptides.

Quantification of murine antigen-specific INFγ levels was carried out by ELISpot (Diaclone, USA) analysis of T cell populations directly *ex vivo*. Ninety-six-well hydrophobic polyvinylidene difluoride membrane-bottomed plates (MAIP S 45; Millipore, USA) were pre-wetted with 70% ethanol. The plates were washed twice with PBS, then coated with anti-INFγ monoclonal antibody at 4°C overnight. After blocking with 2% skimmed milk, plates were washed with PBS, and 100 μL/well of antigen was added in triplicate to appropriate wells. For each assay, a medium-only negative control and a positive control of staphylococcal enterotoxin B (SEB 25 ng/mL) were included. Wells were seeded with 2 x 10^6^ cells/mL in HL-1 medium (supplemented with 1% L-glutamine, 1% penicillin/streptomycin, and 2.5% β-mercaptoethanol) and plates were incubated for 72 hours at 37 °C with 5% CO_2_. The plate contents were then discarded and plates were incubated with PBS/Tween 20 (0.1%) for 10 minutes at 4°C. Plates were then washed twice with PBS/Tween 20 (0.1%) and incubated with biotinylated anti-INFγ monoclonal antibody. Plates were again washed twice with PBS/Tween 20 (0.1%), and then incubated with streptavidin-alkaline phosphatase conjugate. After a wash with PBS/Tween 20 (0.1%), plates were treated with 5-bromo-4-chloro-3-indolyl phosphate and nitro blue tetrazolium (BCIP/NBT) and spot formation was monitored visually. The plate contents were then discarded and plates were washed with water, then air-dried and incubated overnight at 4°C to enhance spot clarity. Spots were counted using an automated ELISpot reader (AID), and results expressed as delta spot-forming cells per 10^6^ cells (ΔSFC/10^6^ which is calculated as SFC/10^6^ of stimulated cells minus SFC/10^6^ of negative control cells). The results were considered positive if the ΔSFC/10^6^ was more than two standard deviations above the negative control.

For assessment of peptide-specific T cell proliferation, murine lymphocytes were resuspended at 3.5×10^6^ cells/mL in supplemented HL-1 media (Lonza, UK) (1% L-glutamine, 1% penicillin/streptomycin, 2.5% β-mercaptoethanol) and 100 μL/well was plated out in triplicate in 96-well Costar tissue culture plates (Corning Incorporated, USA). The cells were stimulated with 100 μL/well of appropriate antigen, positive controls of 5 μg/mL Con A (Sigma-Aldrich, USA) or 25 ng/mL of SEB (Sigma-Aldrich, USA) or negative controls of medium with cells. Plates were incubated at 37°C with 5% CO_2_ for 5 days. Eight hours before harvesting, 1 μCi/well of [^3^H]-thymidine (GE Healthcare, UK) was added. The cells were harvested onto fiberglass filtermats (PerkinElmer, USA) using a Harvester 96 cell harvester (Tomtec, USA) and counted on a Wallac Betaplate scintillation counter (EG&G Instruments, Netherlands). Results were expressed as either delta counts per minute (ΔCPM which is calculated as CPM of stimulated cells minus CPM of negative control cells) or stimulation index (SI which is calculated as CPM of stimulated cells divided by CPM of negative control cells). An SI of ≥ 2.5 was considered to indicate a positive proliferation response.

### PA epitope mapping with human donor PBMC samples

Lymphocytes were isolated from human peripheral blood samples and stimulated as described previously [25]. In brief, sodium-heparinised blood was collected with full informed consent (Ericyes University Ethical Committee) from nine Turkish patients treated for cutaneous anthrax infection within the last eight years and 10 volunteers routinely vaccinated every 12 months for a minimum of five years with the UK AVP vaccine (UK Department of Health under approval by the Convention on Biological Diversity Independent Ethics Committee for the UK Ministry of Defence). Peripheral blood mononuclear cells (PBMC) were isolated from the blood by centrifugation at 800g for 30 minutes in Accuspin tubes (Sigma, UK) cells were then removed from the interface and washed twice in AIM-V serum free media. Cells were counted for viability and resuspended at 2×10^6^ cells/mL.

Human T cell INFγ levels were quantified by ELISpot (Diaclone, France) as previously described [25]. In brief, the peptide library was prepared in a matrix comprising six peptides per pool, so that each peptide occurred in two pools but no peptides occurred together in multiple pools. This allowed the determination of responses to individual peptides. The in-well concentration of each peptide was 25 µg/mL and total peptide concentration per well was 150 µg/mL. After addition of antigen to the wells the plates were frozen at −80 °C until use. Wells were seeded with human PBMCs at 2 x 10^5^ cells/well (range: 1.6 x 10^5^ to 2.1 x 10^5^ cells/well) in AIM-V media (Gibco, UK) and plates were incubated for 72 hours at 37 °C with 5% CO_2_. The plate contents were then discarded and plates were washed with PBS-Tween 20 (0.1%) and incubated with biotinylated anti-INFγ, then washed again before streptavidin-alkaline-phosphatase conjugate was added. After a final wash, plates were developed by addition of BCIP/NBT. Spots were counted using an automated ELISpot reader (AID), and results were expressed as ΔSFC/10^6^. The results were considered positive if the ΔSFC/10^6^ was more than two standard deviations above the negative control and ≥ 50 spots.

### HLA-peptide binding assay

Competitive ELISAs were used to determine the relative binding affinity of PA peptides to HLA-DR molecules as previously described [48, 49]. Briefly, the HLA-DR molecules were immunopurified from homozygous EBV-transformed lymphoblastoid B cell lines by affinity chromatography. The HLA-DR molecules were diluted in HLA binding buffer and incubated for 24 to 72 hours with an appropriate biotinylated reporter peptide, and a serial dilution of the competitor PA peptides. Controls of unlabelled reporter peptides were used as reference peptides to assess the validity of each experiment. 50 μL of HLA binding neutralisation buffer was added to each well and the resulting supernatants were incubated for 2 hours at room temperature in ELISA plates (Nunc, Denmark) previously coated with 10 μg/mL of the monoclonal antibody L243. Bound biotinylated peptide was detected by addition of streptavidin-alkaline phosphatase conjugate (GE Healthcare, France) and 4-methylumbelliferyl phosphate substrate (Sigma-Aldrich, France). Emitted fluorescence was measured at 450 nm post-excitation at 365 nM on a SpectraMax Gemini fluorometer (Molecular Devices, France). The PA peptide concentration that prevented binding of 50% of the labeled peptide (IC_50_) was evaluated, and data expressed as relative binding affinity (ratio of IC_50_ of the PA competitor peptide to the IC_50_ of the reference peptide that binds strongly to the HLA-DR molecule). Sequences of the reference peptides and their IC50 values were as follows: HA 306–318 (PKYVKQNTLKLAT) for DRB1*0101 (4 nM), DRB1*0401 (8 nM), DRB1*1101 (7 nM), YKL (AAYAAAKAAALAA) for DRB1*0701 (3 nM), A3 152–166 (EAEQLRAYLDGTGVE) for DRB1*1501 (48 nM), MT 2–16 (AKTIAYDEEARRGLE) for DRB1*0301 (100 nM), B1 21–36 (TERVRLVTRHIYNREE) for DRB1*1301 (37 nM), DQB45–57 (ADVEVYRAVTPLGPPD) for DQ8 (100 nM) and INS1–15A (FVNQHLAGSHLVEAL) for DQ6 (100nM). Strong binding affinity was defined in this study as a relative activity <100.

## Author Contributions

Conceived and designed the experiments: SA RJI KKC EDW LB SS JHR BM RJB DMA. Performed the experiments: SA RJI KKC HD EDW JHR BM SJM. Analysed the data: SA RJI KKC JHR BM. Contributed reagents/materials/analysis tools: MD GM YO LB SJM TG HD. Wrote the paper: SA RJI RJB DMA. All authors listed have made a substantial, direct, and intellectual contribution to the manuscript and approved it for publication.

## Conflict of Interest Statement

DMA has received payment in a role as scientific consultant to the anthrax vaccine programme at Pfenex Inc. San Diego. The authors declare that this relationship had no role in the study design, data collection and analysis, decision to publish, or preparation of the manuscript.

## Acknowledgements

Julie A. Musson is thanked for assistance with ELISpot assays.

---

**Supporting Figure 1.**
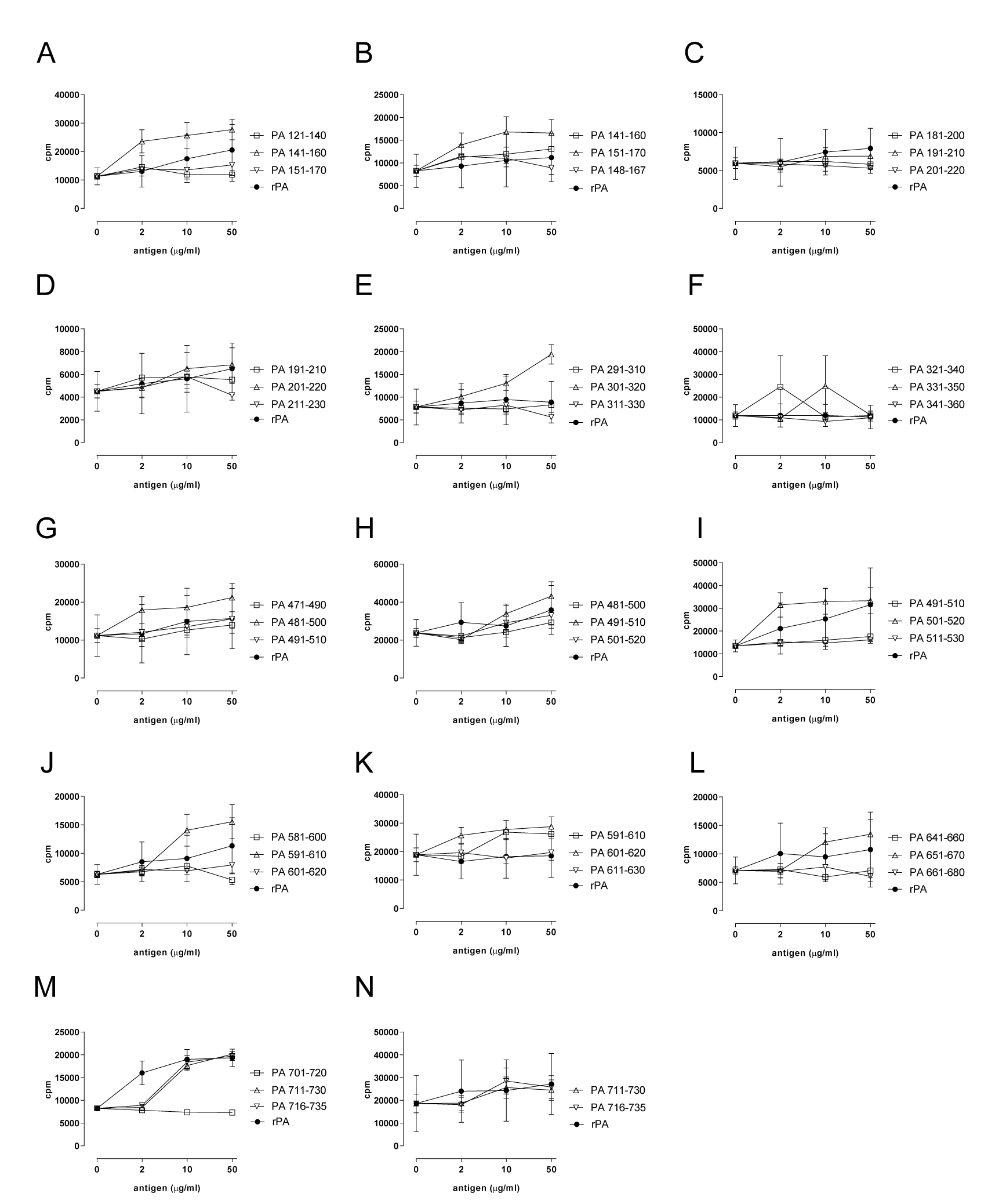
Fine specificity mapping of previously identified HLA-DQ8 restricted T cell epitopes. HLA-DQ8 transgenics were immunised with the previously identified PA peptide in adjuvant. The proliferative responses of draining lymph node cells were measured in response to the indicated concentrations of whole PA protein, domains I-IV of the protein and the immunising and flanking peptides. The responses are shown as the stimulation index calculated as the mean CPM of triplicate wells in the presence of peptide divided by the mean CPM in the absence of antigen. Values twice the mean CPM in the absence of antigen were considered positive responses (n=3 for each data point).

**Supporting Figure 2.**
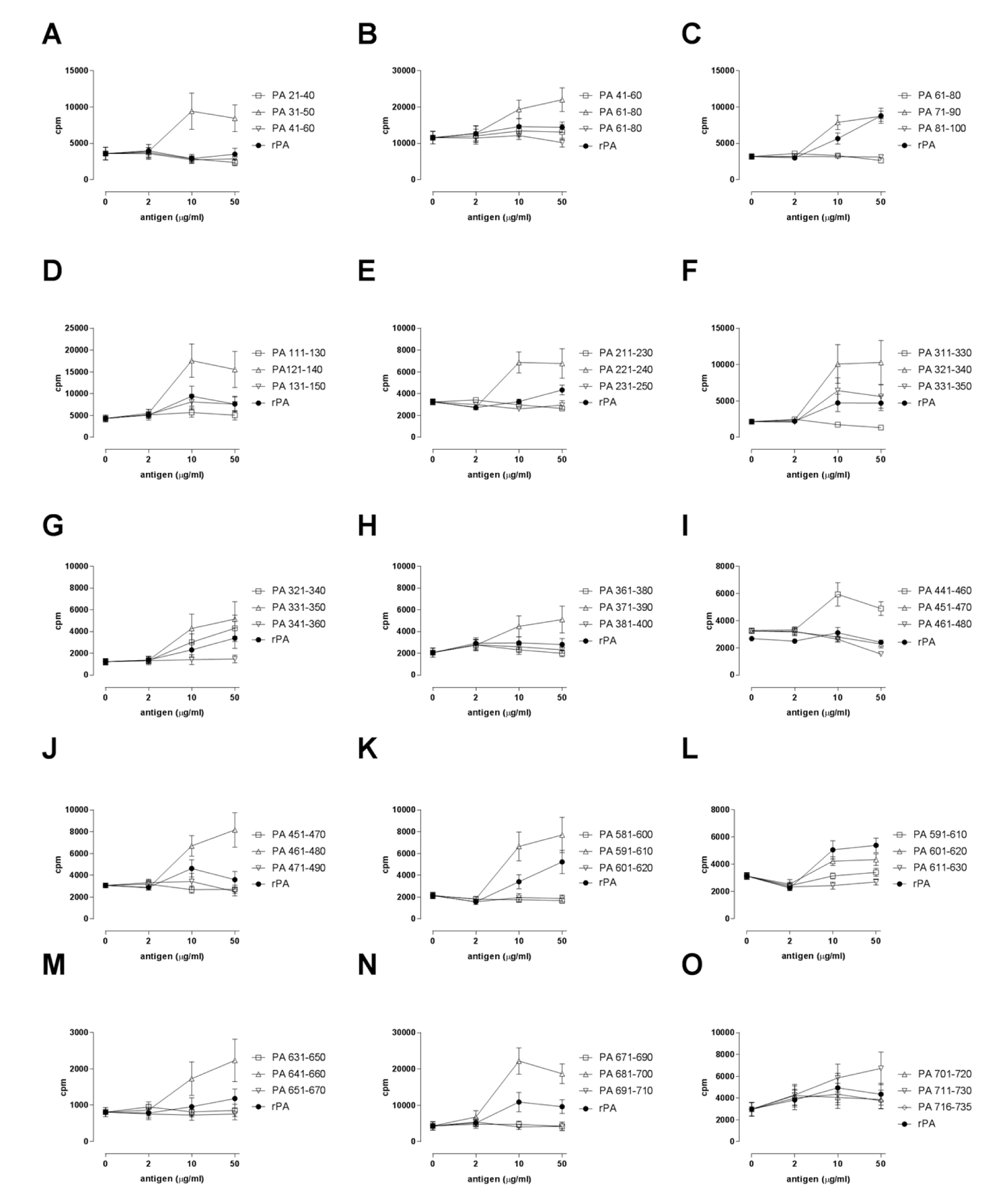
Fine specificity mapping of previously identified HLA-DR4 restricted T cell epitopes. HLA-DR4 transgenics were immunised with the previously identified PA peptide in adjuvant. The proliferative responses of draining lymph node cells were measured in response to the indicated concentrations of whole PA protein, domains I-IV of the protein and the immunising and flanking peptides. The responses are shown as the stimulation index calculated as the mean CPM of triplicate wells in the presence of peptide divided by the mean CPM in the absence of antigen. Values twice the mean CPM in the absence of antigen were considered positive responses (n=3 for each data point).

**Supporting Figure 3.**
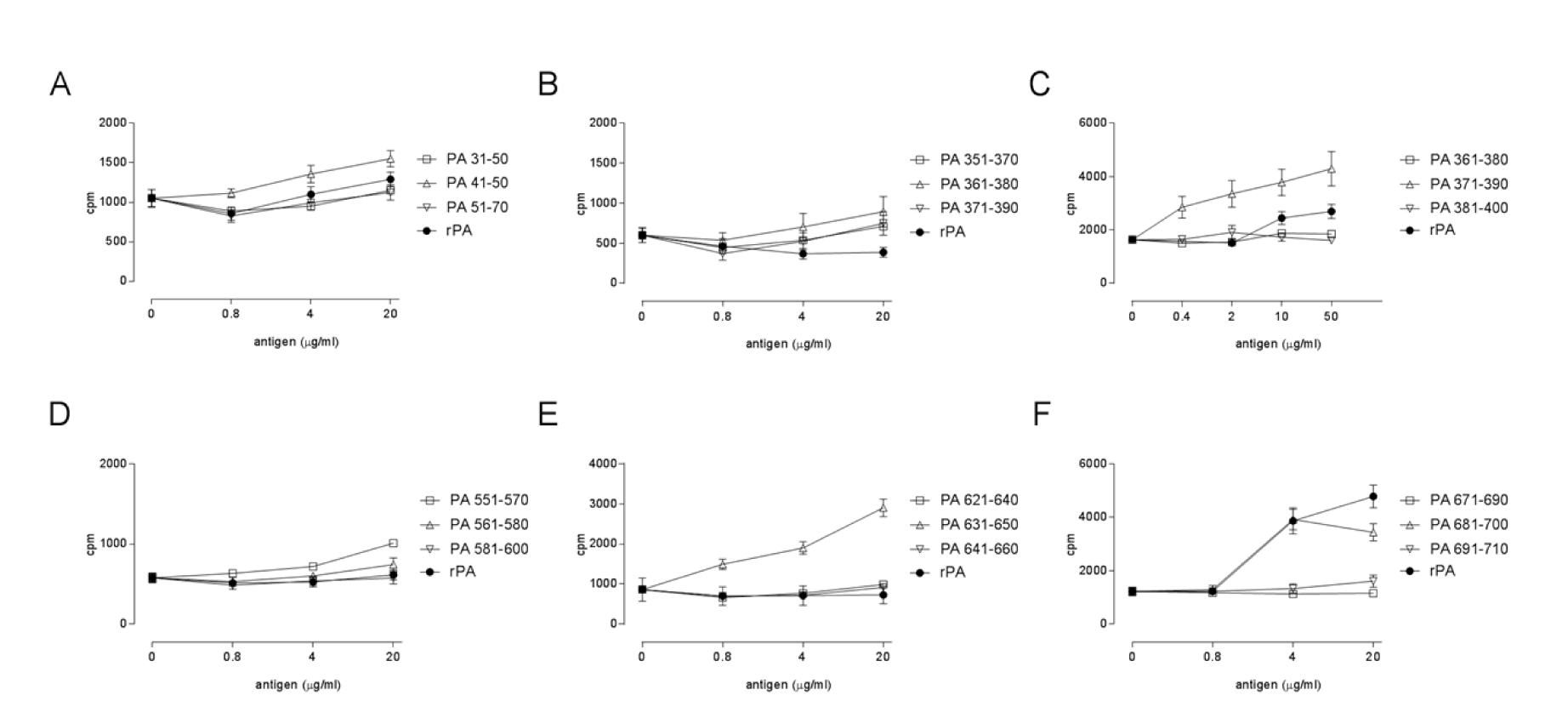
Fine specificity mapping of previously identified HLA-DR1 restricted T cell epitopes. HLA-DR1 transgenics were immunised with the previously identified PA peptide in adjuvant. The proliferative responses of draining lymph node cells were measured in response to the indicated concentrations of whole PA protein, domains I-IV of the protein and the immunising and flanking peptides. The responses are shown as the stimulation index calculated as the mean CPM of triplicate wells in the presence of peptide divided by the mean CPM in the absence of antigen. Values twice the mean CPM in the absence of antigen were considered positive responses (n=3 for each data point).

**Supporting Table 1.**
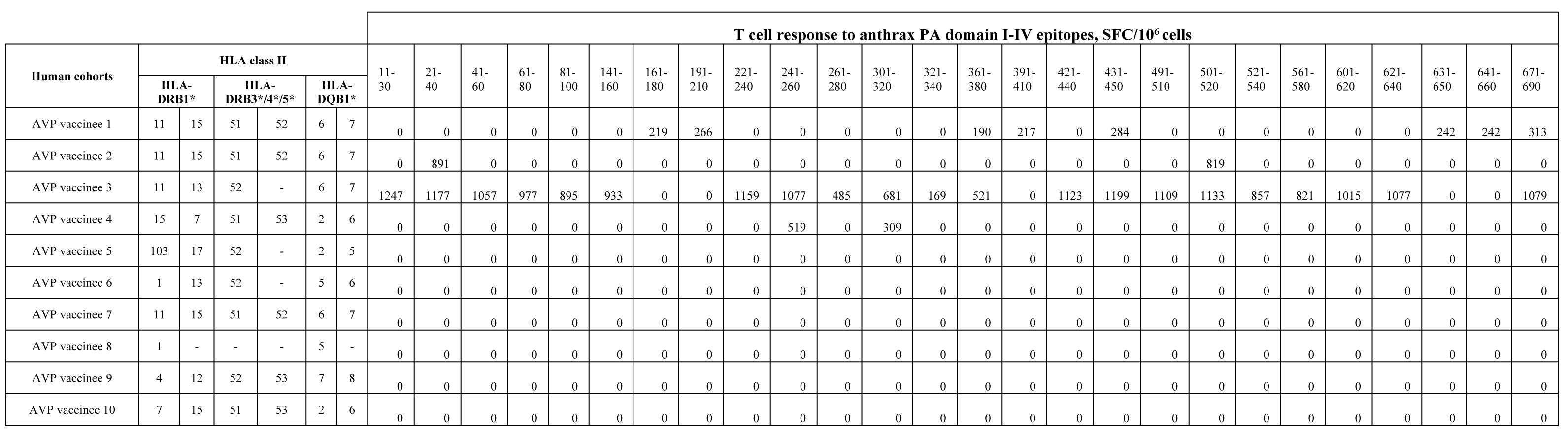
CD4+ T cell responses to B. anthracis PA epitopes in AVP vaccinees.

**Supporting Table 2.** CD4+ T cell responses to B. anthracis PA epitopes in anthrax-recovered patients.

|  |  |  |  |  |  |  | T cell response to anthrax PA domain I-IV epitopes, SFC/10<sup>6</sup> cells |  |  |  |  |  |  |  |  |  |  |  |  |  |  |
| --- | --- | --- | --- | --- | --- | --- | --- | --- | --- | --- | --- | --- | --- | --- | --- | --- | --- | --- | --- | --- | --- |
| Human cohorts | HLA class II |  |  |  |  |  | 1-20 | 11-30 | 21-40 | 31-50 | 41-60 | 51-70 | 61-80 | 71-90 | 81-100 | 91-110 | 101-120 | 121-140 | 131-150 | 141-160 | 151-170 |
|  | HLA-DRB1* |  | HLA-DRB3*/4*/5* |  | HLA-DQB1* |  |  |  |  |  |  |  |  |  |  |  |  |  |  |  |  |
| Infected donor 1 | 11 | 13 | 52 | - | 6 | 7 | 0 | 0 | 0 | 0 | 0 | 0 | 0 | 0 | 0 | 0 | 0 | 0 | 0 | 0 | 0 |
| Infected donor 2 | 4 | - | 53 | - | 8 | - | 299 | 330 | 301 | 263 | 224 | 0 | 203 | 0 | 0 | 0 | 0 | 0 | 304 | 299 | 417 |
| Infected donor 3 | 4 | 14 | 52 | 53 | 5 | 8 | 0 | 0 | 0 | 0 | 0 | 0 | 0 | 486 | 0 | 454 | 0 | 0 | 0 | 0 | 0 |
| Infected donor 4 | 15 | - | 51 | - | 6 | - | 0 | 0 | 0 | 0 | 0 | 0 | 0 | 0 | 0 | 0 | 0 | 0 | 0 | 0 | 0 |
| Infected donor 5 | 8 | 11 | 52 | - | 7 | - | 0 | 0 | 0 | 0 | 0 | 0 | 0 | 0 | 0 | 0 | 0 | 0 | 0 | 0 | 977 |
| Infected donor 6 | 11 | 13 | 52 | - | 6 | 7 | 0 | 0 | 0 | 0 | 0 | 0 | 0 | 0 | 0 | 0 | 0 | 581 | 0 | 0 | 0 |
| Infected donor 7 | 4 | 14 | 52 | 53 | 5 | - | 0 | 0 | 1017 | 0 | 971 | 723 | 1193 | 957 | 1225 | 949 | 793 | 451 | 0 | 505 | 691 |
| Infected donor 8 | 1 | 16 | 51 | - | 5 | - | 0 | 0 | 0 | 0 | 759 | 784 | 0 | 0 | 0 | 0 | 0 | 0 | 0 | 0 | 0 |
| Infected donor 9 | 15 | 13 | 51 | 52 | 6 | - | 0 | 0 | 0 | 0 | 0 | 0 | 0 | 0 | 0 | 0 | 0 | 0 | 0 | 0 | 0 |

| Human cohorts | T cell response to anthrax PA domain I-IV epitopes, SFC/10<sup>6</sup> cells |  |  |  |  |  |  |  |  |  |  |  |  |  |  |  |  |
| --- | --- | --- | --- | --- | --- | --- | --- | --- | --- | --- | --- | --- | --- | --- | --- | --- | --- |
|  | 148-167 | 168-187 | 161-180 | 171-190 | 181-200 | 191-210 | 211-230 | 221-240 | 231-250 | 241-260 | 261-280 | 271-290 | 281-300 | 301-320 | 311-330 | 321-340 | 331-350 |
| Infected donor 1 | 0 | 0 | 0 | 0 | 0 | 0 | 0 | 0 | 0 | 0 | 0 | 0 | 0 | 0 | 0 | 0 | 0 |
| Infected donor 2 | 337 | 383 | 349 | 268 | 0 | 229 | 229 | 246 | 0 | 304 | 0 | 0 | 0 | 176 | 0 | 0 | 0 |
| Infected donor 3 | 0 | 0 | 0 | 0 | 0 | 0 | 0 | 0 | 0 | 0 | 0 | 0 | 0 | 0 | 0 | 0 | 0 |
| Infected donor 4 | 0 | 0 | 0 | 0 | 0 | 0 | 0 | 0 | 0 | 0 | 0 | 0 | 0 | 0 | 0 | 0 | 0 |
| Infected donor 5 | 0 | 0 | 0 | 0 | 0 | 0 | 0 | 0 | 0 | 0 | 0 | 0 | 0 | 0 | 0 | 0 | 0 |
| Infected donor 6 | 0 | 0 | 0 | 0 | 0 | 0 | 0 | 0 | 0 | 0 | 0 | 0 | 641 | 0 | 0 | 0 | 477 |
| Infected donor 7 | 1013 | 919 | 1251 | 915 | 911 | 921 | 0 | 0 | 1117 | 711 | 757 | 1063 | 759 | 649 | 799 | 805 | 685 |
| Infected donor 8 | 0 | 813 | 0 | 0 | 0 | 0 | 0 | 0 | 0 | 0 | 723 | 0 | 0 | 0 | 0 | 212 | 0 |
| Infected donor 9 | 0 | 263 | 0 | 0 | 0 | 0 | 0 | 0 | 0 | 0 | 0 | 0 | 0 | 0 | 0 | 0 | 0 |

|  | T cell response to anthrax PA domain I-IV epitopes, SFC/106 cells |  |  |  |  |  |  |  |  |  |  |  |  |  |  |  |  |
| --- | --- | --- | --- | --- | --- | --- | --- | --- | --- | --- | --- | --- | --- | --- | --- | --- | --- |
| Human cohorts | 341-360 | 351-370 | 361-380 | 371-390 | 381-400 | 391-410 | 401-420 | 411-430 | 421-440 | 431-450 | 441-460 | 451-470 | 461-480 | 471-490 | 481-500 | 491-510 | 501-520 |
| Infected donor 1 | 0 | 0 | 0 | 0 | 0 | 0 | 0 | 0 | 0 | 0 | 0 | 0 | 0 | 0 | 0 | 0 | 0 |
| Infected donor 2 | 0 | 263 | 325 | 304 | 0 | 263 | 215 | 268 | 323 | 299 | 0 | 232 | 179 | 222 | 0 | 325 | 289 |
| Infected donor 3 | 0 | 0 | 0 | 0 | 0 | 0 | 0 | 0 | 0 | 0 | 0 | 0 | 0 | 0 | 0 | 0 | 0 |
| Infected donor 4 | 0 | 0 | 0 | 0 | 0 | 0 | 0 | 0 | 0 | 0 | 0 | 0 | 0 | 0 | 0 | 0 | 0 |
| Infected donor 5 | 0 | 0 | 0 | 0 | 0 | 0 | 0 | 0 | 0 | 0 | 0 | 0 | 0 | 0 | 0 | 0 | 0 |
| Infected donor 6 | 0 | 0 | 0 | 0 | 0 | 0 | 0 | 0 | 0 | 599 | 801 | 0 | 0 | 0 | 577 | 0 | 0 |
| Infected donor 7 | 891 | 0 | 747 | 735 | 855 | 1199 | 0 | 0 | 859 | 1039 | 0 | 899 | 0 | 0 | 807 | 0 | 661 |
| Infected donor 8 | 0 | 0 | 0 | 0 | 0 | 0 | 0 | 0 | 0 | 0 | 0 | 0 | 0 | 0 | 0 | 0 | 0 |
| Infected donor 9 | 0 | 0 | 0 | 0 | 0 | 0 | 0 | 0 | 0 | 0 | 0 | 0 | 0 | 0 | 0 | 0 | 0 |
| Human cohorts | 511-530 | 521-540 | 531-550 | 541-560 | 551-570 | 561-580 | 571-590 | 581-600 | 591-610 | 601-620 | 611-630 | 621-640 | 631-650 | 641-660 | 651-670 | 661-680 | 671-690 |
| Infected donor 1 | 0 | 0 | 0 | 0 | 0 | 0 | 0 | 0 | 0 | 0 | 0 | 0 | 0 | 0 | 0 | 0 | 0 |
| Infected donor 2 | 241 | 234 | 0 | 306 | 0 | 0 | 0 | 0 | 232 | 234 | 301 | 311 | 414 | 357 | 270 | 256 | 222 |
| Infected donor 3 | 0 | 0 | 0 | 0 | 0 | 0 | 396 | 0 | 0 | 0 | 0 | 0 | 0 | 0 | 0 | 0 | 0 |
| Infected donor 4 | 0 | 0 | 0 | 0 | 0 | 0 | 0 | 0 | 0 | 0 | 0 | 0 | 0 | 0 | 0 | 0 | 0 |
| Infected donor 5 | 1061 | 0 | 0 | 0 | 0 | 0 | 0 | 0 | 0 | 0 | 0 | 0 | 0 | 0 | 0 | 0 | 0 |
| Infected donor 6 | 0 | 0 | 603 | 0 | 0 | 0 | 495 | 779 | 0 | 0 | 0 | 0 | 539 | 0 | 0 | 0 | 545 |
| Infected donor 7 | 419 | 815 | 995 | 1391 | 881 | 1085 | 983 | 619 | 0 | 0 | 0 | 621 | 1043 | 0 | 893 | 705 | 993 |
| Infected donor 8 | 0 | 617 | 846 | 0 | 0 | 0 | 0 | 0 | 0 | 0 | 0 | 0 | 0 | 0 | 823 | 0 | 0 |
| Infected donor 9 | 0 | 0 | 0 | 0 | 596 | 0 | 0 | 0 | 0 | 0 | 0 | 0 | 0 | 0 | 474 | 0 | 0 |

| Human cohorts | T cell response to anthrax PA domain I-IV epitopes, SFC/106 cells |  |  |
| --- | --- | --- | --- |
|  | 681-700 | 691-710 | 716-735 |
| Infected donor 1 | 0 | 0 | 0 |
| Infected donor 2 | 0 | 210 | 0 |
| Infected donor 3 | 532 | 0 | 0 |
| Infected donor 4 | 0 | 0 | 0 |
| Infected donor 5 | 0 | 0 | 0 |
| Infected donor 6 | 905 | 0 | 0 |
| Infected donor 7 | 929 | 1121 | 869 |
| Infected donor 8 | 0 | 0 | 0 |
| Infected donor 9 | 0 | 0 | 0 |

**Supporting Table 3.** Differential susceptibility of HLA class II transgenic mice to anthrax infection.

| HLA | *B. anthracis*<br>STI<br>challenge<br>dose (CFU) | Number of<br>mice<br>challenged | Number<br>of<br>challenge<br>survivors | Mean time<br>to death<br>(days) | Bacterial load in<br>spleens within<br>observation period<br>post-infection mean<br>CFU/spleen (± SEM) | Bacterial load<br>in spleens of<br>survivors at<br>day 20 mean<br>CFU/spleen<br>(± SEM) | Estimated<br>LD<sub>50</sub> (CFU) |
| --- | --- | --- | --- | --- | --- | --- | --- |
| C57Bl6 | 10<sup>5</sup> | 10 | 4 | 4.5 (±0.22) | 1.0x10<sup>3</sup> (±0.32x10<sup>3</sup>)<br>(n=6) | 91 (±18) | 10<sup>5</sup> |
| DQ6 | 10<sup>5</sup> | 8 | 8 | - | - | 255 (±39) | >10<sup>5</sup> |
| DR4 | 10<sup>5</sup> | 10 | 8 | 7 (±3) | 1.27x10<sup>3</sup><br>(n=1) | 22 (±13) | >10<sup>5</sup> |
| DR15 | 10<sup>5</sup> | 9 | 5 | 5.75<br>(±0.48) | 0.85x10<sup>3</sup> (±0.21 x10<sup>3</sup>)<br>(n=4) | 67 (±32) | 10<sup>5</sup> |
| DQ8 | 10<sup>6</sup> | 10 | 8 | 6 | 2.17x10<sup>3</sup> (±0.67x10<sup>3</sup>)<br>(n=2) | 896 (±263) | >10<sup>6</sup> |
| DR1 | 10<sup>6</sup> | 10 | 10 | - | - | 411 (±93) | >10<sup>6</sup> |

## References

1. Goel AK. Anthrax: A disease of biowarfare and public health importance. World J Clin Cases. 2015;3(1):20–33. Epub 2015/01/23. doi: 10.12998/wjcc.v3.i1.20. PubMed PMID: 25610847; PubMed Central PMCID: PMCPMC4295216.

2. Dixon TC, Meselson M, Guillemin J, Hanna PC. Anthrax. N Engl J Med. 1999;341(11):815–26. Epub 1999/09/09. doi: 10.1056/NEJM199909093411107. PubMed PMID: 10477781.

3. Green MS, LeDuc J, Cohen D, Franz DR. Confronting the threat of bioterrorism: realities, challenges, and defensive strategies. Lancet Infect Dis. 2019;19(1):e2–e13. Epub 2018/10/21. doi: 10.1016/S1473-3099(18)30298-6. PubMed PMID: 30340981.

4. Abbara A, Brooks T, Taylor GP, Nolan M, Donaldson H, Manikon M, et al. Lessons for control of heroin-associated anthrax in Europe from 2009-2010 outbreak case studies, London, UK. Emerg Infect Dis. 2014;20(7):1115–22. Epub 2014/06/25. doi: 10.3201/eid2007.131764. PubMed PMID: 24959910; PubMed Central PMCID: PMCPMC4073855.

5. Revich BA, Podolnaya MA. Thawing of permafrost may disturb historic cattle burial grounds in East Siberia. Glob Health Action. 2011;4. Epub 2011/11/25. doi: 10.3402/gha.v4i0.8482. PubMed PMID: 22114567; PubMed Central PMCID: PMCPMC3222928.

6. Baillie LW, Fowler K, Turnbull PC. Human immune responses to the UK human anthrax vaccine. J Appl Microbiol. 1999;87(2):306–8. Epub 1999/09/04. PubMed PMID: 10475977.

7. Chitlaru T, Altboum Z, Reuveny S, Shafferman A. Progress and novel strategies in vaccine development and treatment of anthrax. Immunol Rev. 2011;239(1):221–36. Epub 2011/01/05. doi: 10.1111/j.1600-065X.2010.00969.x. PubMed PMID: 21198675.

8. Enstone JE, Wale MC, Nguyen-Van-Tam JS, Pearson JC. Adverse medical events in British service personnel following anthrax vaccination. Vaccine. 2003;21(13- 14):1348–54. Epub 2003/03/05. PubMed PMID: 12615429.

9. Brey RN. Molecular basis for improved anthrax vaccines. Adv Drug Deliv Rev. 2005;57(9):1266–92. Epub 2005/06/07. doi: 10.1016/j.addr.2005.01.028. PubMed PMID: 15935874.

10. Hopkins RJ, Kalsi G, Montalvo-Lugo VM, Sharma M, Wu Y, Muse DD, et al. Randomized, double-blind, active-controlled study evaluating the safety and immunogenicity of three vaccination schedules and two dose levels of AV7909 vaccine for anthrax post-exposure prophylaxis in healthy adults. Vaccine. 2016;34(18):2096–105. Epub 2016/03/17. doi: 10.1016/j.vaccine.2016.03.006. PubMed PMID: 26979136; PubMed Central PMCID: PMCPMC4839983.

11. Baillie LW. Past, imminent and future human medical countermeasures for anthrax. J Appl Microbiol. 2006;101(3):594–606. Epub 2006/08/16. doi: 10.1111/j.1365-2672.2006.03112.x. PubMed PMID: 16907809.

12. Brown BK, Cox J, Gillis A, VanCott TC, Marovich M, Milazzo M, et al. Phase I study of safety and immunogenicity of an Escherichia coli-derived recombinant protective antigen (rPA) vaccine to prevent anthrax in adults. PLoS One. 2010;5(11):e13849. Epub 2010/11/17. doi: 10.1371/journal.pone.0013849. PubMed PMID: 21079762; PubMed Central PMCID: PMCPMC2974626.

13. Campbell JD, Clement KH, Wasserman SS, Donegan S, Chrisley L, Kotloff KL. Safety, reactogenicity and immunogenicity of a recombinant protective antigen anthrax vaccine given to healthy adults. Hum Vaccin. 2007;3(5):205–11. Epub 2007/09/21. PubMed PMID: 17881903.

14. Gorse GJ, Keitel W, Keyserling H, Taylor DN, Lock M, Alves K, et al. Immunogenicity and tolerance of ascending doses of a recombinant protective antigen (rPA102) anthrax vaccine: a randomized, double-blinded, controlled, multicenter trial. Vaccine. 2006;24(33-34):5950–9. Epub 2006/06/27. doi: 10.1016/j.vaccine.2006.05.044. PubMed PMID: 16797805.

15. Hewetson JF, Little SF, Ivins BE, Johnson WM, Pittman PR, Brown JE, et al. An in vivo passive protection assay for the evaluation of immunity in AVA-vaccinated individuals. Vaccine. 2008;26(33):4262–6. Epub 2008/07/01. doi: 10.1016/j.vaccine.2008.05.068. PubMed PMID: 18586363.

16. Smith K, Crowe SR, Garman L, Guthridge CJ, Muther JJ, McKee E, et al. Human monoclonal antibodies generated following vaccination with AVA provide neutralization by blocking furin cleavage but not by preventing oligomerization. Vaccine. 2012;30(28):4276–83. Epub 2012/03/20. doi: 10.1016/j.vaccine.2012.03.002. PubMed PMID: 22425791; PubMed Central PMCID: PMCPMC3367042.

17. Reuveny S, White MD, Adar YY, Kafri Y, Altboum Z, Gozes Y, et al. Search for correlates of protective immunity conferred by anthrax vaccine. Infect Immun. 2001;69(5):2888–93. Epub 2001/04/09. doi: 10.1128/IAI.69.5.2888-2893.2001. PubMed PMID: 11292703; PubMed Central PMCID: PMCPMC98239.

18. Crowe SR, Ash LL, Engler RJ, Ballard JD, Harley JB, Farris AD, et al. Select human anthrax protective antigen epitope-specific antibodies provide protection from lethal toxin challenge. J Infect Dis. 2010;202(2):251–60. Epub 2010/06/11. doi: 10.1086/653495. PubMed PMID: 20533877; PubMed Central PMCID: PMCPMC2891133.

19. Quinn CP, Sabourin CL, Niemuth NA, Li H, Semenova VA, Rudge TL, et al. A three-dose intramuscular injection schedule of anthrax vaccine adsorbed generates sustained humoral and cellular immune responses to protective antigen and provides long-term protection against inhalation anthrax in rhesus macaques. Clin Vaccine Immunol. 2012;19(11):1730–45. Epub 2012/08/31. doi: 10.1128/CVI.00324-12. PubMed PMID: 22933399; PubMed Central PMCID: PMCPMC3491539.

20. McBride BW, Mogg A, Telfer JL, Lever MS, Miller J, Turnbull PC, et al. Protective efficacy of a recombinant protective antigen against Bacillus anthracis challenge and assessment of immunological markers. Vaccine. 1998;16(8):810–7. Epub 1998/06/17. PubMed PMID: 9627938.

21. Williamson ED, Beedham RJ, Bennett AM, Perkins SD, Miller J, Baillie LW. Presentation of protective antigen to the mouse immune system: immune sequelae. J Appl Microbiol. 1999;87(2):315–7. Epub 1999/09/04. PubMed PMID: 10475979.

22. Zhang Y, Qiu J, Zhou Y, Farhangfar F, Hester J, Lin AY, et al. Plasmid-based vaccination with candidate anthrax vaccine antigens induces durable type 1 and type 2 T-helper immune responses. Vaccine. 2008;26(5):614–22. Epub 2008/01/02. doi: 10.1016/j.vaccine.2007.11.072. PubMed PMID: 18166249.

23. Doolan DL, Freilich DA, Brice GT, Burgess TH, Berzins MP, Bull RL, et al. The US capitol bioterrorism anthrax exposures: clinical epidemiological and immunological characteristics. J Infect Dis. 2007;195(2):174–84. Epub 2006/12/28. doi: 10.1086/510312. PubMed PMID: 17191162.

24. Glomski IJ, Corre JP, Mock M, Goossens PL. Cutting Edge: IFN-gamma-producing CD4 T lymphocytes mediate spore-induced immunity to capsulated Bacillus anthracis. J Immunol. 2007;178(5):2646–50. Epub 2007/02/22. PubMed PMID: 17312104.

25. Ingram RJ, Metan G, Maillere B, Doganay M, Ozkul Y, Kim LU, et al. Natural exposure to cutaneous anthrax gives long-lasting T cell immunity encompassing infection-specific epitopes. J Immunol. 2010;184(7):3814–21. Epub 2010/03/09. doi: 10.4049/jimmunol.0901581. PubMed PMID: 20208010.

26. Ascough S, Ingram RJ, Chu KK, Reynolds CJ, Musson JA, Doganay M, et al. Anthrax lethal factor as an immune target in humans and transgenic mice and the impact of HLA polymorphism on CD4+ T cell immunity. PLoS Pathog. 2014;10(5):e1004085. Epub 2014/05/03. doi: 10.1371/journal.ppat.1004085. PubMed PMID: 24788397; PubMed Central PMCID: PMCPMC4006929.

27. Ascough S, Ingram RJ, Abarra A, Holmes AJ, Maillere B, Altmann DM, et al. Injectional anthrax infection due to heroin use induces strong immunological memory. J Infect. 2014;68(2):200–3. Epub 2014/02/12. doi: 10.1016/j.jinf.2013.10.007. PubMed PMID: 24513100; PubMed Central PMCID: PMCPMC4150029.

28. Ascough S, Ingram RJ, Chu KK, Musson JA, Moore SJ, Gallagher T, et al. CD4+ T Cells Targeting Dominant and Cryptic Epitopes from Bacillus anthracis Lethal Factor. Front Microbiol. 2015;6:1506. Epub 2016/01/19. doi: 10.3389/fmicb.2015.01506. PubMed PMID: 26779161; PubMed Central PMCID: PMCPMC4700811.

29. Ingram RJ, Ascough S, Reynolds CJ, Metan G, Doganay M, Baillie L, et al. Natural cutaneous anthrax infection, but not vaccination, induces a CD4(+) T cell response involving diverse cytokines. Cell Biosci. 2015;5:20. Epub 2015/06/16. doi: 10.1186/s13578-015-0011-4. PubMed PMID: 26075052; PubMed Central PMCID: PMCPMC4464127.

30. Ascough S, Altmann DM. Anthrax in injecting drug users: the need for increased vigilance in the clinic. Expert Rev Anti Infect Ther. 2015;13(6):681–4. Epub 2015/04/02. doi: 10.1586/14787210.2015.1032255. PubMed PMID: 25831413.

31. Laws TR, Kuchuloria T, Chitadze N, Little SF, Webster WM, Debes AK, et al. A Comparison of the Adaptive Immune Response between Recovered Anthrax Patients and Individuals Receiving Three Different Anthrax Vaccines. PLoS One. 2016;11(3):e0148713. Epub 2016/03/24. doi: 10.1371/journal.pone.0148713. PubMed PMID: 27007118; PubMed Central PMCID: PMCPMC4805272.

32. Altmann DM. Host immunity to Bacillus anthracis lethal factor and other immunogens: implications for vaccine design. Expert Rev Vaccines. 2015;14(3):429–34. Epub 2014/11/18. doi: 10.1586/14760584.2015.981533. PubMed PMID: 25400140.

33. Williamson ED, Hodgson I, Walker NJ, Topping AW, Duchars MG, Mott JM, et al. Immunogenicity of recombinant protective antigen and efficacy against aerosol challenge with anthrax. Infect Immun. 2005;73(9):5978–87. Epub 2005/08/23. doi: 10.1128/IAI.73.9.5978-5987.2005. PubMed PMID: 16113318; PubMed Central PMCID: PMCPMC1231098.

34. Chen L, Schiffer JM, Dalton S, Sabourin CL, Niemuth NA, Plikaytis BD, et al. Comprehensive analysis and selection of anthrax vaccine adsorbed immune correlates of protection in rhesus macaques. Clin Vaccine Immunol. 2014;21(11):1512–20. Epub 2014/09/05. doi: 10.1128/CVI.00469-14. PubMed PMID: 25185577; PubMed Central PMCID: PMCPMC4248764.

35. Glomski IJ, Piris-Gimenez A, Huerre M, Mock M, Goossens PL. Primary involvement of pharynx and peyer’s patch in inhalational and intestinal anthrax. PLoS Pathog. 2007;3(6):e76. Epub 2007/06/05. doi: 10.1371/journal.ppat.0030076. PubMed PMID: 17542645; PubMed Central PMCID: PMCPMC1885272.

36. Ingram R, Baillie L. It’s in the genes! Human genetic diversity and the response to anthrax vaccines. Expert Rev Vaccines. 2012;11(6):633–5. Epub 2012/08/10. doi: 10.1586/erv.12.41. PubMed PMID: 22873120.

37. Garman L, Dumas EK, Kurella S, Hunt JJ, Crowe SR, Nguyen ML, et al. MHC class II and non-MHC class II genes differentially influence humoral immunity to Bacillus anthracis lethal factor and protective antigen. Toxins (Basel). 2012;4(12):1451–67. Epub 2013/01/25. PubMed PMID: 23342680; PubMed Central PMCID: PMCPMC3528256.

38. Pajewski NM, Parker SD, Poland GA, Ovsyannikova IG, Song W, Zhang K, et al. The role of HLA-DR-DQ haplotypes in variable antibody responses to anthrax vaccine adsorbed. Genes Immun. 2011;12(6):457–65. Epub 2011/03/04. doi: 10.1038/gene.2011.15. PubMed PMID: 21368772; PubMed Central PMCID: PMCPMC3165112.

39. Li ZK, Nie JJ, Li J, Zhuang H. The effect of HLA on immunological response to hepatitis B vaccine in healthy people: a meta-analysis. Vaccine. 2013;31(40):4355–61. Epub 2013/07/28. doi: 10.1016/j.vaccine.2013.06.108. PubMed PMID: 23887040.

40. Posteraro B, Pastorino R, Di Giannantonio P, Ianuale C, Amore R, Ricciardi W, et al. The link between genetic variation and variability in vaccine responses: systematic review and meta-analyses. Vaccine. 2014;32(15):1661–9. Epub 2014/02/12. doi: 10.1016/j.vaccine.2014.01.057. PubMed PMID: 24513009.

41. Agrawal A, Lingappa J, Leppla SH, Agrawal S, Jabbar A, Quinn C, et al. Impairment of dendritic cells and adaptive immunity by anthrax lethal toxin. Nature. 2003;424(6946):329–34. Epub 2003/07/18. doi: 10.1038/nature01794. PubMed PMID: 12867985.

42. Phillips-Conroy JE, Hildebolt CF, Altmann J, Jolly CJ, Muruthi P. Periodontal health in free-ranging baboons of Ethiopia and Kenya. Am J Phys Anthropol. 1993;90(3):359–71. Epub 1993/03/01. doi: 10.1002/ajpa.1330900310. PubMed PMID: 8460659.

43. Nojima M, Ihara H, Kyo M, Hashimoto M, Ito K, Kunikata S, et al. The significant effect of HLA-DRB1 matching on acute rejection in kidney transplants. Transpl Int. 1996;9 Suppl 1:S11–5. Epub 1996/01/01. PubMed PMID: 8959780.

44. Ellmerich S, Takacs K, Mycko M, Waldner H, Wahid F, Boyton RJ, et al. Disease-related epitope spread in a humanized T cell receptor transgenic model of multiple sclerosis. Eur J Immunol. 2004;34(7):1839–48. Epub 2004/06/24. doi: 10.1002/eji.200324044. PubMed PMID: 15214032.

45. Ellmerich S, Mycko M, Takacs K, Waldner H, Wahid FN, Boyton RJ, et al. High incidence of spontaneous disease in an HLA-DR15 and TCR transgenic multiple sclerosis model. J Immunol. 2005;174(4):1938–46. Epub 2005/02/09. PubMed PMID: 15699121.

46. Boyton RJ, Lohmann T, Londei M, Kalbacher H, Halder T, Frater AJ, et al. Glutamic acid decarboxylase T lymphocyte responses associated with susceptibility or resistance to type I diabetes: analysis in disease discordant human twins, non-obese diabetic mice and HLA-DQ transgenic mice. Int Immunol. 1998;10(12):1765–76. Epub 1999/01/14. PubMed PMID: 9885897.

47. Flick-Smith HC, Walker NJ, Gibson P, Bullifent H, Hayward S, Miller J, et al. A recombinant carboxy-terminal domain of the protective antigen of Bacillus anthracis protects mice against anthrax infection. Infect Immun. 2002;70(3):1653–6. Epub 2002/02/21. PubMed PMID: 11854261; PubMed Central PMCID: PMCPMC127760.

48. Texier C, Pouvelle S, Busson M, Herve M, Charron D, Menez A, et al. HLA-DR restricted peptide candidates for bee venom immunotherapy. J Immunol. 2000;164(6):3177–84. Epub 2000/03/08. PubMed PMID: 10706708.

49. Pancre V, Georges B, Angyalosi G, Castelli F, Delanoye A, Delacre M, et al. Novel promiscuous HLA-DQ HIV Nef peptide that induces IFN-gamma-producing memory CD4+ T cells. Clin Exp Immunol. 2002;129(3):429–37. Epub 2002/08/29. doi: 10.1046/j.1365-2249.2002.01934.x. PubMed PMID: 12197883; PubMed Central PMCID: PMCPMC1906467.

50. Petosa C, Collier RJ, Klimpel KR, Leppla SH, Liddington RC. Crystal structure of the anthrax toxin protective antigen. Nature. 1997;385(6619):833–8. Epub 1997/02/27. doi: 10.1038/385833a0. PubMed PMID: 9039918.

